# Of Rats and Men, a Translational Model to Understand Vancomycin Pharmacokinetic/Toxicodynamic relationships

**DOI:** 10.1101/2021.04.22.437975

**Authors:** Marc H. Scheetz, Gwendolyn Pais, Thomas P. Lodise, Steven Y.C. Tong, Joshua S. Davis, J. Nicholas O’Donnell, Jiajun Liu, Michael Neely, Walter C. Prozialeck, Peter C. Lamar, N.Jim Rhodes, Thomas Holland, Sean N. Avedissian

## Abstract

**Background:** Vancomycin is a first line antibiotic for many common infectious diseases and is the most commonly prescribed antibiotic in the United States hospital setting. Vancomycin is also well known to cause kidney injury; two recent prospective studies have identified that increasing vancomycin area under the concentration curve predicts vancomycin induced kidney injury (VIKI). However, outside of clinical trials, it is unclear if pre-clinical data can quantitatively describe VIKI in patients.

**Methods:** Data were simultaneously analyzed from a pre-clinical rat model and two prospective clinical studies. Logged vancomycin area under the concentration curve (AUC) data for rats (n=48) and patients from PROVIDE (n=263) and CAMERA2 (n=291) were included. VIKI was defined as urinary KIM-1 concentrations ≥9.42 ng/mL in the rat and according to KDIGO stage 1 kidney injury for all human patients. Multiple generalized linear models were explored, and the order of magnitude was calculated between the probability of acute kidney injury (AKI) from the average obtained in the clinical studies (i.e. CAMERA2 and PROVIDE) and the rat for 0.1 increments in Log10AUC bounded common concentrations obtained in the therapeutic range (i.e. ~200 −800 mg*24h/L).

**Results:** A logit link model best fit the data. When calculating the multiplicative factors between the studies therapeutic range AUCs, the rat was an average 2.7 to 4.2 times more sensitive to AKI between AUCs of 199.5 (i.e. log 10 AUC=2.3) and 794.3 mg*h/L (i.e. log 10 AUC=2.9), respectively.

**Conclusions:** A pre-clinical rat model was quantitatively linked to toxicity data from two large human studies. The rat is an attractive pre-clinical model to explore exposure toxicity relationships with vancomycin. External validation is required.

## Introduction

As a first line antibiotic in consensus guidelines for serious and life-threatening infections,(14, 17, 19, 44, 49, 50) vancomycin is the single most commonly prescribed antibiotic in the United States hospital setting.(12, 18, 25, 36, 37) In addition to common use, vancomycin is a drug well known to increase the risk of AKI.(38) A prospective study identified an excess 10% attributable AKI risk to vancomycin when compared to linezolid for treatment of methicillin-resistant *Staphylococcus aureus (*MRSA) pneumonia.(56) Based on 36.5 million hospital stays in the US annually(55) and vancomycin prevalence use of ~100 days of therapy/1000 patient days,(12, 18) even conservative estimates suggest that vancomycin causes kidney injury in over 300,000 people annually.

Vancomycin induced kidney injury (VIKI) is a concentration-driven process, with area under the concentration curve (AUC) and maximum concentrations best correlating with the extent of kidney damage.(8–10) Studies indicate that rates of VIKI were approximately 5-7% when troughs were maintained between 5-10 mg/L as the standard of practice.(27, 43, 46) With the publication of the 2009 consensus vancomycin guidelines that promoted more aggressive dosing (i.e. troughs of 15-20 mg/L) for serious MRSA infections such as pneumonia (5, 45), studies indicate that VIKI is now considerably higher, with rates upwards of 40% depending on the patient population, without any improvements in effectiveness.(46) In an effort to minimize VIKI while maintaining comparable effectiveness, the revised 2020 vancomycin guidelines tempered targeted exposures and recommended AUC-rather than trough-based dosing, with a daily AUC target of 400-600 mg*h/L for serious MRSA infections.(41)

Support to transition from trough-only to AUC-guided dosing and monitoring was based, in part, on two recent prospective analyses that evaluated the efficacy and safety of vancomycin against MRSA bacteremia. In both the PROVIDE(23) and CAMERA2 (20, 51) studies, VIKI was found to increase as a function of the daily AUC, especially among patients with daily AUCs in excess of 600 mg*h/L. However, patients with vancomycin exposures within the newly recommended daily AUC therapeutic range of 400-600 mg*h/L were also found to be at increased risk of VIKI. (20, 23) The exact magnitude of AKI attributable to vancomycin in these studies is unclear as they lacked a control group who did not receive vancomycin and many patients who received vancomycin had other risk factors for AKI. However, these data suggest that over half the cases were likely attributable to vancomycin.(47) Given the high frequency of vancomycin use and considerable potential for VIKI associated with maintaining daily AUCs within the newly recommended range of 400-600 mg*h/L, there is a critical need to identify vancomycin exposure profiles that minimize VIKI in clinical practice. Ideally, it would be preferred to identify optimal vancomycin dosing and monitoring practices that minimize VIKI in a clinical trial. However, the costs and time associated with execution of well-designed studies in patients greatly limits the ability to generate timely results. Therefore, a pre-clinical model is sorely needed that is reflective of the vancomycin exposure-VIKI experience in patients as it will surmount many of the aforementioned barriers associated with completion of clinical studies and provide quantitative information in a shorter timeframe. In particular, the availability of a reliable and accurate pre-clinical VIKI model will identify vancomycin dosing schemes associated with the lowest risk of VIKI, determine if ameliorating agents can minimize the risk of VIKI with vancomycin exposures required for efficacy, and assess populations in which vancomycin should be avoided (e.g., those receiving other nephrotoxic agents). Our group has utilized a pre-clinical rat model that describes the risk of VIKI as a function of the intensity of vancomycin exposure(9, 31, 34, 39). While our model demonstrates that there is a clear relationship between vancomycin AUC and VIKI in rats, we have yet to determine if our model bridges to humans. To this end, we sought to evaluate the relationship between vancomycin AUC and VIKI from our translational rat model and the clinical studies, PROVIDE and CAMERA2(23, 51).

## Methods

### Data Sources

#### Animal data

The relationship between vancomycin exposure and VIKI were obtained from our previously published rat study.(10) In brief, this pharmacokinetic/toxicodynamic PK/TD study (IACUC; Protocol #2295) was conducted at Midwestern University in Downers Grove, IL in compliance with the National Institutes of Health Guide for the Care and Use of Laboratory Animals.[(1)]. In this study, male Sprague-Dawley rats (n=48, approximately 8-10 weeks old, mean weight 310g) received intravenous saline (controls) or intravenous vancomycin (150 mg/kg/day to 400 mg/kg/day as once or twice daily dosing for a period of 24 hours). The dosing range was selected previous studies (10, 15, 30, 52) to ensure coverage of the clinical allometric range. For example, the clinical kidney injury threshold of ≥ 4 grams/day in a 70-kg patient (i.e., 57 mg/kg/day in humans) scales allometrically to 350 mg/kg in the rat. (4, 21) Plasma was sampled for vancomycin assay (completed by LC-MS/MS) with up to 8 samples per rat over the course of the study. Twenty-four-hour urine was collected and assayed for kidney injury molecule 1 (KIM-1). PK exposures were obtained from each individual animal, with area under the concentration curve for the 24-hour period calculated in Pmetrics for R.(29) The 90th percentile effective concentration, EC_90_, (i.e. vancomycin concentration required to achieve 90^th^ percent maximum of KIM-1) was experimentally calculated from the fitted Hill model. Results were utilized as reported in the parent publication (10) with the exception that two errors were found in the calculation of KIM-1 concentrations. These errors did not affect any of the published summary results or exposure response fits reported in the parent publication.

### Clinical Data

#### CAMERA2

CAMERA2 was an open-label, international, pragmatic, randomized clinical trial performed at 27 hospitals between 2015 and 2018. The trial enrolled 352 hospitalized adults with MRSA bacteremia. Patients randomly received either 1) vancomycin or daptomycin plus an anti-staphylococcal β-lactam (intravenous flucloxacillin, cloxacillin, or cefazolin) (n = 174) or vancomycin or daptomycin alone (n = 178).(51) Among these patients, 291 patients had their individual vancomycin exposures [i.e. area under the concentration curve (AUC)] estimated with a best-fit Bayesian PK model in a post-hoc analysis.(20) AUCs were calculated over 24 hour periods, and day 2 AUC best correlated with acute kidney injury outcomes described by modified-Kidney Disease Improving Global Outcomes (m-KDIGO) criteria (6, 20, 51) as well as risk, injury, failure, loss, and end-stage kidney disease (RIFLE) criteria(16). AUC24-48h cut-points for prediction of m-KDIGO ≥1, m-KDIGO ≥2, and m-KDIGO =3 were 470.1, 496.1, and 525.5.(20)

### PROVIDE

PROVIDE was a prospective, observational study performed at 14 hospitals between 2014 and 2015.(23) The study enrolled 265 hospitalized adults treated with vancomycin for their MRSA bacteremia. Patients received vancomycin therapeutic drug monitoring, and day 2 AUCs were estimated from a Bayesian maximal a posteriori probability procedure approach. Patients were followed for treatment success and acute kidney injury. Kidney injury was defined by RIFLE criteria and vancomycin-induced nephrotoxicity (VINT) definition in the vancomycin consensus guideline statement (42). Outcomes were further classified using a desirability of outcome ranking (DOOR) analysis. Efficacy was defined by 30-day mortality and lack of persistent bacteremia. The five categories were: death, survival with treatment failure and AKI, survival with treatment failure and no AKI, survival with treatment success and AKI, and finally survival with treatment success and no AKI. Patients with a day 2 AUC ≥793 had higher rates of AKI and VIN vs to those with an AUC ≤343. Patients in the 2 lowest AUC exposure quintiles (i.e. AUC ≤515), had the best global outcome (i.e. survival with treatment success and no AKI) with rates ≥71% vs. ≤61% respectively.

### Definition of AKI event

In the rat studies, AKI events were evaluated with urinary KIM-1 (Figure 1). Urinary KIM-1 was chosen as the biomarker for linking as it has been demonstrated as the most sensitive and specific biomarker for predicting histopathologic damage in the rat.(33) For the rat, the vancomycin AUC EC_90_ was 482 mg*h/L, corresponding to a urinary Log_2_ KIM equal to 2^3.236^=9.42 ng/mL. The Log_2_ transformation of KIM is prudent as KIM-1 elevations vs. control have been suggested to follow fold changes, and control animals display concentrations less than 2^1^=2 ng/mL. Thus, the threshold for injury was experimentally defined as approximately two doubling fold changes, Figure 2.(10) Importantly, this value also classified rat kidney injury well by histopathology. In this analysis, histopathology scores ranged from 0-3 using the Predictive Safety Testing Consortium criteria from a blinded veterinary pathologist,(48) and control animals universally scored 0 or 1. The KIM-1 threshold correctly classified histopathology scores of 2 with a sensitivity of 81.8% and 3 with a sensitivity of 100%. In a secondary and exploratory analysis, histopathology scores of 2 or greater were used to define AKI for the rat in this study.

**Figure 1.**
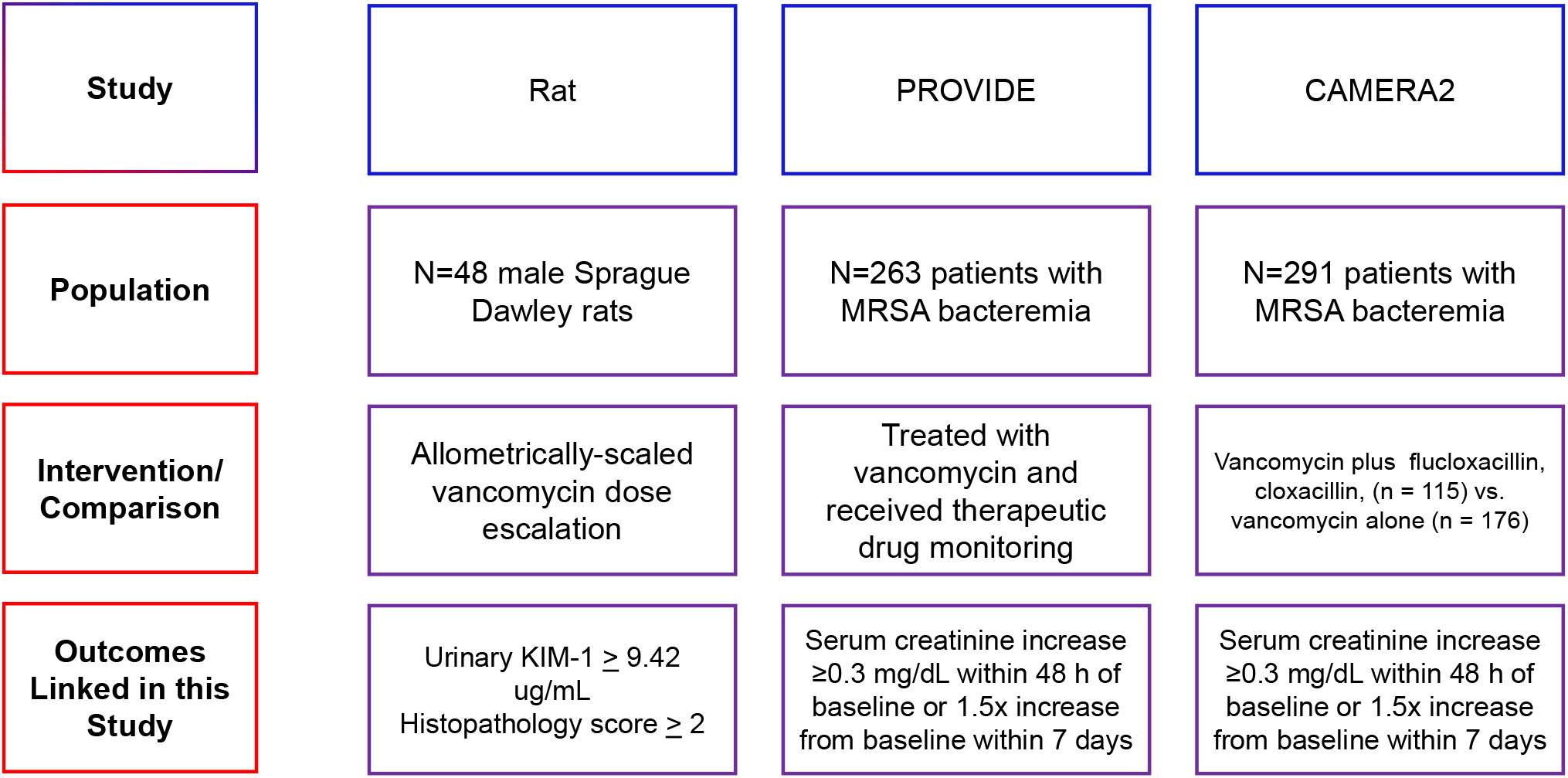
Overview of the primary data utilized in this manuscript.

**Figure 2.**
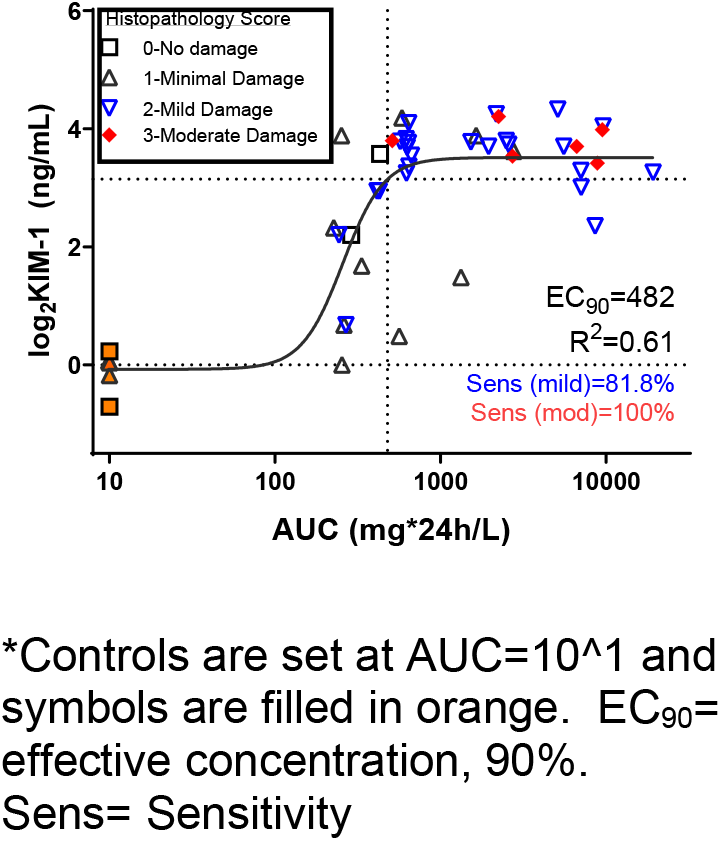
Relationship between vancomycin AUC and urinary log_2_KIM

For the two clinical analyses, a common kidney injury endpoint was used; KDIGO stage 1 criteria defined VIKI (20, 23, 51), and all patients that could be classified according to this endpoint were included (Figure 1). The KDIGO endpoint was chosen (i.e. serum creatinine [SCr] absolute increase of ≥0.3 mg/dL within 48 hours of baseline or 1.5x SCr increase from baseline within 7 days) to define VIKI as data shows that absolute changes in creatinine (vs. relative percentage changes) are more reliable for detection of renal insult when compared to measures that rely on relative changes (24, 35, 53). In total, n=48, n=263, and n=291 subjects were included from the rat study,(10) PROVIDE (23), and CAMERA2 (20, 51) respectively. Five rats were controls and received no vancomycin.

### Statistical methods

Statistical analysis was performed in Stata/IC version 16.1 (College Station, TX) unless otherwise specified. All data were pooled, and subjects were categorized according to their parent study. VIKI was dichotomous in each dataset, and the probability of VIKI served as the endpoint. For the animal data, the day 1 AUC served as the primary exposure variable while the day 2 AUC was the primary vancomycin exposure variable in the clinical studies (23) (20). Day 2 was utilized for the human data as this has been the most consistent predictor of AKI from the prospective clinical studies. Multiple generalized linear models with maximum likelihood optimization were explored, including binomial family with logit and probit link with VIKI as the dichotomous outcome and vancomycin AUC as the primary predictor variable. The exact study (i.e. rat, PROVIDE, CAMERA2) was included as a categorical covariate in the model. Interactions were checked between AUCs and study group. Model fits were compared with the Akaike information criterion (AIC) scores. Probabilities for AKI from each of the studies was predicted for incremental AUCs (mg/L*hr) as a log_10_ function between 0 and 6 with an increment of 0.1. Margins were calculated to estimate individual probabilities of AKI for each subject according to each study and across predicted log_10_ AUC from the final fitted model. The order of magnitude was calculated between the probability of AKI from the average obtained in the clinical studies (i.e. CAMERA2 and PROVIDE) and the rat for 0.1 increments in Log_10_AUCs in the therapeutic range (i.e. ~200 - 800 mg*24h/L).

### Human and animal assurances

Clinical data were collected under parent IRBs (20, 23). Analysis of de-identified data was obtained under data use agreements and classified as not-human-subjects research by the Midwestern University IRB. Rat work was conducted under Institutional Animal Care and Use Committee (IACUC) Protocol #2295.

## Results

AUCs from the rats receiving vancomycin ranged from 226.74 to 19,239 mg*h/L, with a median of 643.1 and an interquartile range of 427.7 to 2769.4. AUCs in PROVIDE and CAMERA2 ranged from 94.3 to 1755 mg*h/L and 159.3 to 1211.4 mg*h/L, respectively with a median of 578.1 and 398.7 and IQRs of 436.2 to 548.1 and 300.9 to 526.6 mg*h/L, respectively. A total of 63% of rats experienced kidney injury based on KIM-1 threshold and 67.4% experienced kidney injury based on secondary outcome of a histopathology threshold. AKI rates for PROVIDE and CAMERA2 patients were 17.5%, and 17.2%, respectively.

Model fits for the binomial family with logit link (AIC 550.6) were slightly better than with probit link (AIC 551.6). Thus, logit models were used for application. Interactions were not present between study group and subject AUCs (p≥ 0.45). Log_10_AUC, as a continuous function, predicted VIKI across all three datasets well. Results from the logit link model, transformed to odds ratios for the primary outcome, can be found in Table 1. Probabilities of AKI were a function of AUC (P<0.001) as well as individual study (P<0.001 for both), Figure 3. Predicted probabilities did not significantly differ whether rat AKI was classified by urinary KIM-1 value (Figure 3) or histopathologic cut-point (Figure S1). The 95% confidence intervals for CAMERA2 and PROVIDE completely overlapped for the range of AUCs studied (data not shown). When calculating the multiplicative factors between the studies for therapeutic AUCs, the rat was an average 2.1 to 3.1 times more sensitive to AKI across AUCs of 199.5 (i.e. log 10 AUC=2.3) to 794.3 mg*h/L (i.e. log 10 AUC=2.9), respectively (Table 2).

**Table 1.** Independent odds of kidney injury with the rat model as the referent group.

|  | Odds Ratio | P value | 95% CI |  |
| --- | --- | --- | --- | --- |
| log 10 AUC | 8.97 | P<0.001 | 3.23 | 24.87 |
| Study |  |  |  |  |
| Rat | 1 [referent] |  |  |  |
| PROVIDE | 0.15 | P<0.001 | 0.68 | 0.34 |
| CAMERA2 | 0.20 | P<0.001 | 0.09 | 0.45 |

**Figure 3.**
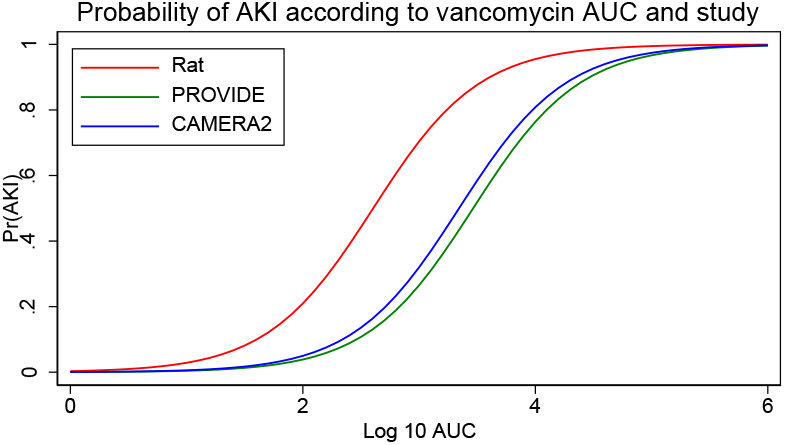
Probability of AKI for each subject and across a span of vancomycin AUCs as calculated from the final fitted model.

**Table 2.**
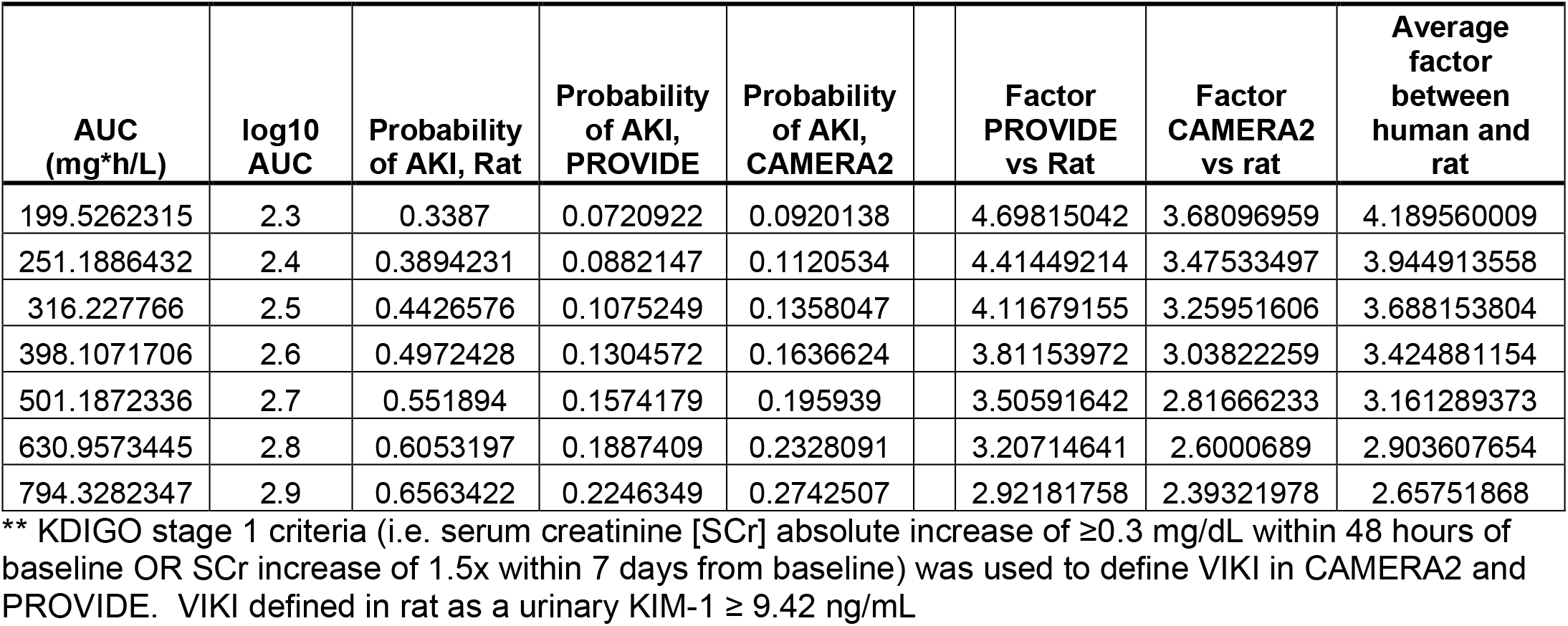
Probabilities of AKI across commonly achieve AUCs within the vancomycin therapeutic range and linking factor between the studies.

## Discussion

Overall, we found that VIKI increased as a function of the daily Log_10_ AUC in a predictable fashion across the rat and two human studies. Quantiatively, the rat model was 2.1 to 3.1 times more sensitive in detecting VIKI across therapeutic AUCs observed in PROVIDE and CAMERA2. The quantitative link across the studies indicates that the rat model can be used to reliably forecast the expected rate of VIKI at a given AUC value in adult patients receiving vancomycin. Functionally, this can be interpreted with the plotted probabilities (e.g. Figure 3). That is, for any given dose plotted on the x-axis, the expected probability of event (i.e. the rat achieving urinary KIM-1 concentrations ≥9.42 ng/mL or histopathology ≥ 2 in a 24 hour model and humans achieving serum creatinine absolute increase of ≥0.3 mg/dL within 48 hours of baseline or 1.5x SCr increase from baseline within 7 days). Importantly, confirmation that our rat model is highly translatable to humans has important implications for clinical practice as our linked model provides an efficient way to identify optimal candidate vancomycin dosing strategies that minimize VIKI for potential use into clinical practice. Clinical trials will always be the gold standard for assessing exposure-response relationships but they are expensive (costs are a median of $41,000 per enrolled patient (28), are generally designed to answer a single question, and take many years to complete. Directly relevant, CAMERA2 cost ~ $US 2M and took 4 years to complete whereas PROVIDE cost ~5 million dollars and required 3 years to complete. Translatable pre-clinical models can also be used to provide immediate insights into questions such as: “Can VIKI be minimized by altering the concentration-time curve?; Can VIKI be ameliorated with co-administration of a prophylactic or rescue agent?; Does the therapeutic window change when common co-nephrotoxins are given?” All of these questions are commonly faced clinically, yet each question requires a clinical trial. With limited resources, pre-clinical models such as the one evaluated in this study provide a screening mechanism to ensure that only the most promising ideas are evaluated in a clinical trial.

The demonstrable link between the rat and human studies are biologically plausible. The rat is a well-developed model for acute kidney injury, and newer biomarkers such as KIM-1 are highly relevant as they are shared between humans and rats.(2, 3) KIM-1 is a specific marker of histopathologic proximal tubule injury in VIKI (11, 30) as well as is qualified by the FDA for drug induced acute kidney injury in both human and rat drug studies.(7) Our findings were isometric depending on whether we used a KIM-1 or a histopathological cut-point. For the VIKI model, we favored using KIM-1 as urinary samples are easy to obtain longitudinally (and do not require animal sacrifice). Further, KIM-1 is very sensitive for prediction of histopathologic damage (Figure 1) which is still considered by some as the gold standard for translational toxicological studies.

Several points should be noted when evaluating the findings. The rat model relied on day 1 estimates of vancomycin exposure in 24 hour experiments and used KIM-1 to define VIKI. We believe it was appropriate to use the AUC:KIM-1 data from the 24-hour rat studies in the exposure response analyses and link them to day 2 AUC:SCr VIKI endpoints in the human studies. We have observed in our model that KIM-1 increases on day 1 with vancomycin treatment and plateaus for several days in our more prolonged experiments (34). Thus, one day experiments appear sufficient to define the injury profile. In the translational pre-clinical model, the goal is to obtain a marker that is not in the causal pathway for injury and thus measure a predictor rather than an intermediate surrogate for toxicity that has already occurred (e.g. vancomycin AUC increases because glomerular filtration has already decreased).

In CAMERA2 and PROVIDE, the vancomycin exposure-VIKI response curve was explained by the day 2 AUC(20, 23). We also believe it was appropriate to examine the day 2 AUC in patients relative to day 1 as it is more indicative of near steady-state conditions and the maintenance vancomycin regimen patients received during the early course of vancomycin therapy. In clinical patients, day 1 concentration profiles are also potentially more variable than day 2, as varying renal function is more prevalent during the initial period when patients are being managed for sepsis. Day 2 was also selected for the clinical data because this is when vancomycin concentration-time data would be most likely obtained (22, 26, 56). Ultimately, the best approach for future human trials will be to see if early biomarkers such as KIM-1 can predict clinical toxicity before the more standard clinical markers such as serum creatinine. This would enable clinicians to change therapy prior to more substantial damage, and once data for KIM-1 become available from human trials, the rat model can be re-calibrated.

In humans, SCr is the common clinical biomarker for defining acute kidney injury (6), but it is known to be a delayed indicator of renal injury and decline in renal function(57). Because of renal reserve, SCr only increases after a substantial amount of damage has occurred to the nephrons. It is estimated that greater than 40% reductions are needed in CrCL in order to observe creatinine changes within reasonable time frames such as within 48 hours (40, 54). Based on the amount of pre-existing kidney damage, the SCr may take 24–36 hours to rise after a definite renal insult (13, 32, 54). Therefore, we examined VIKI events downstream from the initial exposure as the injury process likely started several days prior to SCr elevation in human studies. In contrast, urinary KIM-1 rises are detectable within 9 hours of insult (52) and are sensitive and specific for proximal tubule damage specifically with VIKI (33). Thus, KIM-1 is an ideal maker for VIKI, allowing for the most proximal linking between antecedent vancomycin exposure and damage. This toxicologic model can be envisioned similar to other clinical PK/PD analyses in which the achieving of early exposure targets is linked to outcomes like clinical response at test of cure or 28-day mortality. *In vivo* or *in vitro* models utilized for prediction of outcomes are simplifications (by necessity) of the human condition and outcomes.

In PROVIDE, the unadjusted and adjusted risk ratio between AUC and VIKI endpoints were nearly identical, suggesting that the observed results were accurate on both the population and patient level and not modified significantly by covariates. In CAMERA2, receipt of flucloxacillin resulted in more kidney injury. In a sensitivity analysis in which we separated patients by their receipt of flucloxacillin, it was observed that patients who received flucloxacillin had rates of VIKI according to AUC that were similar to the whole population PROVIDE dataset (Figure S2), yet overall relationships from our primary analysis were highly explanatory (Figure2) so we more conservatively included all patients from CAMERA2 (as opposed to restricting analyses only to patients with flucloxacillin).

This study has several limitations. First, these animal studies have been performed in one laboratory and appear to have good internal validity; however, external validation is required to confirm if others are able to replicate similar exposure response curves in the rat and determine if individual models require calibration. Second, we utilized all available data meeting inclusion criteria requirements from the clinical studies and did not make any adjustments for patient covariates (e.g. co-nephrotoxins) in our primary analysis. A sensitivity analysis identified that patients that received flucloxacillin in CAMERA2 demonstrated similar VIKI rates for matched AUCs in the patient population from PROVIDE. The current analyses demonstrate that linking rat data to human data at the population level is possible. Future work will need to test covariates in the rat and recalibrate a link with the stratified human data.

## Conclusion

We have demonstrated that a rat model links to kidney outcomes from the clinical studies, PROVIDE and CAMERA2. The rat is a useful model that has the potential to provide quantitative information on the shift of the vancomycin toxicity curve in humans. Rat models can be applied to focus on distinct questions of interest (such as combinatorial therapy) and can serve as an initial assessment before clinical trials are conducted, thus improving understanding in the setting of no clinical data and clinical data that suffers from confounding relationships. External validation and replication are needed to verify the translational nature of our animal model.

---

**Figure S1.**
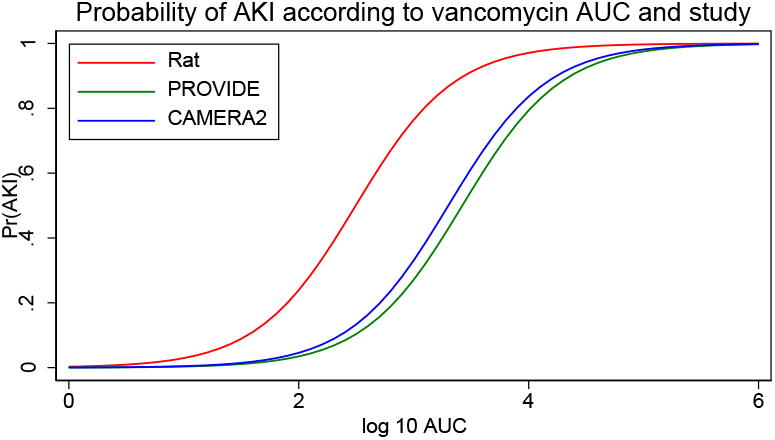
Probability of AKI for each subject and across a span of vancomycin AUCs as calculated from the final fitted model. Rat AKI was defined by histopathology.

**Figure S2.**
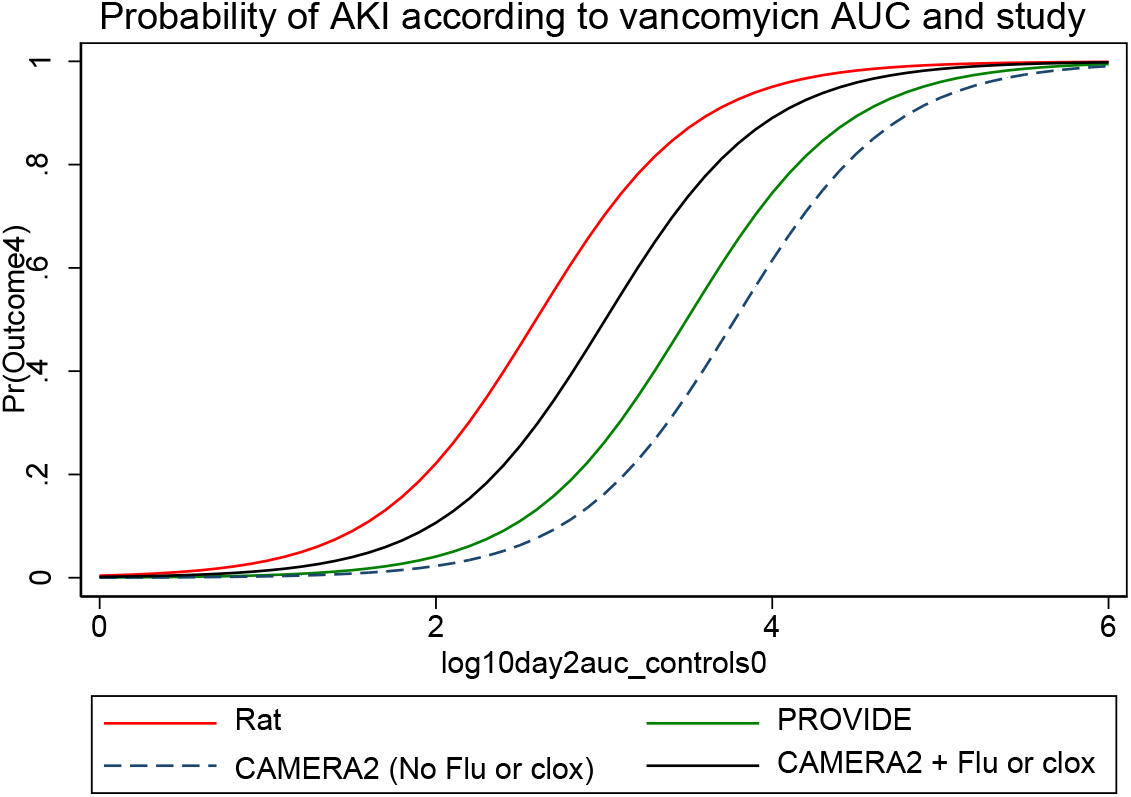
VIKI probability by study, where CAMERA2 is divided into patients that received flucloxacillin and those that did not.

